# Biological traits and threats interact to drive extinctions in a simulation study

**DOI:** 10.1101/2020.10.18.344473

**Authors:** Filipe Chichorro, Luís Correia, Pedro Cardoso

## Abstract

How a particular threat influences extinction risk may depend on biological traits. Empirical studies relating threats and traits are needed, but data are scarce, making simulations useful. We implemented an eco-evolutionary model to analyse how five threat types influence the extinction risk of virtual organisms differing in body size, maturity age, fecundity, and dispersal ability. Results show that direct killing mostly affected slow-living and low dispersal organisms. Habitat loss and fragmentation both affected larger and less fecund organisms, but drove contrasting responses according to dispersal ability. Habitat degradation and the introduction of invasive competitors had similar effects, mostly affecting large, fast-living, and highly fecund organisms. Many of the reported results confirm previous studies, while others were never tested, creating new hypotheses for future empirical work.

**Statement of authorship:** FC, LC and PC designed the study, FC implemented the model and ran the statistical analyses. FC and PC wrote the first draft, and all authors contributed substantially to further revisions.

## Introduction

Species are going extinct at unprecedented rates in human history (Vos *et al*. 2015). Understanding what drives species to extinction is crucial if further extinctions are to be minimized approaching natural, pre-human, levels. Yet, the mechanisms leading to extinction, both intrinsic (species traits) and extrinsic (environmental threats), and the way they interact remain obscure (Chichorro *et al*. 2020).

Much progress has been made recently, with over 150 studies trying to identify similarities in traits between threatened species in the last two decades (Chichorro *et al*. 2019). These studies were both comparative and correlative in the sense that they compared traits of threatened species with those of non-threatened species, often using the IUCN red list categories (IUCN 2020) as proxies of extinction risk (Purvis *et al*. 2000; Cardillo *et al*. 2005; González-Suárez *et al*. 2013; Marco *et al*. 2015).

A recent study (Chichorro *et al*. 2020) tested, for the first time, multiple traits related to extinction risk across a large part of the Tree of Life and the entire world, minimizing functional, phylogenetic, and spatial biases of previous studies. Using data from close to 900 species of vertebrates, invertebrates, and plants, it was found that five traits, namely habitat breadth, fecundity, dispersal ability, altitudinal range, and degree of human influence were universal predictors of extinction risk. The response of five other traits, i.e. body size, offspring size, generation length, microhabitat, and change in the degree of human influence across the species’ range was dependent on the taxon, often on the threat type affecting each taxon. It showed large promise for future studies that the mechanisms driving their similarities and differences across taxa and threat types could be elucidated.

Previous correlative studies indicate that external factors interact in important ways with the relation between intrinsic traits and extinction risk (Olden *et al*. 2007; González-Suárez *et al*. 2013; Murray *et al*. 2014). Habitat destruction, pollution, invasive species, overexploitation, and climate change (Secretariat of the Convention on Biological Diversity 2014) all drive extinctions, and depending on the driver, some traits might be particularly relevant in a species trajectory towards extinction.

In a “ statistically ideal” world, we would have full data on species traits, external drivers, and potential mechanisms linking these to extinction. However, as models become more complex, the amount of data required to test hypotheses increases. Additionally, because observations are rarely fully independent (dispersal ability might be connected with body size, fecundity with offspring size, etc.), it is often difficult to partition this complexity in independent components.

Conceptual models have been suggested to complement correlative models in order to better understand mechanistic drivers of diversity patterns (Murray *et al*. 2014). Within these, individual-based models (or agent-based models, henceforth referred to as ABMs) are increasingly used to aid understanding of the functioning of species and ecosystems (DeAngelis & Grimm 2014). Their power derives from the possibility of replicating patterns found at the population and community levels from phenomena occurring at the individual level (the level at which the basic deciding biological entity exists). Despite difficulties in specifying and implementing these models, today tools exist that facilitate both specification (Grimm *et al*. 2006, 2010) and implementation (Wilensky 1999) of ABMs.

In this study, we develop a conceptual ABM to evaluate the influence of different threats, namely direct killing, habitat loss, habitat fragmentation, habitat degradation, and invaders, on the risk of extinction of species presenting differing body size, maturity age, fecundity, and dispersal ability. The landscape includes organisms competing for resources, which they use to invest in survival, growth, and reproduction. After creating virtual species adapted to given simulated worlds, we apply the threats and observe how the average trait values of populations respond to each threat. These results from *in silico* experiments are related to real-world knowledge on different organisms and reveal the mechanisms behind species extinction. Additionally, we hypothesize mechanisms to be tested in the future, which can guide data collection, and can be confirmed or disproved as more empirical data become available.

## Materials and methods

### The model

The purpose of the model was to analyse the shift in mean trait values of an evolving population of organisms competing for resources, when an external threat event was introduced into the virtual world where they lived (Fig. 1). We implemented it as an ABM that simulated a landscape where organisms moved in the environment in order to obtain the required resources to survive and reproduce. At initialization, organisms presented a random set of traits. The population then converged towards normally distributed trait values around a stable mean. Each stable population (after convergence) was then treated as an experimental unit. We subjected each experimental unit to a range of threats at several intensities. As the population converged to a new set of trait value distributions, we registered the variation in mean trait values occurring under each threat type and intensity. The ABM model was developed in Netlogo 6.1.1 (Wilensky 1999). A full description of the model following the Overview Design Concepts and Details (ODD) protocol (Grimm *et al*. 2006, 2010) is available in Supplementary Materials S1.

**Figure 1:**
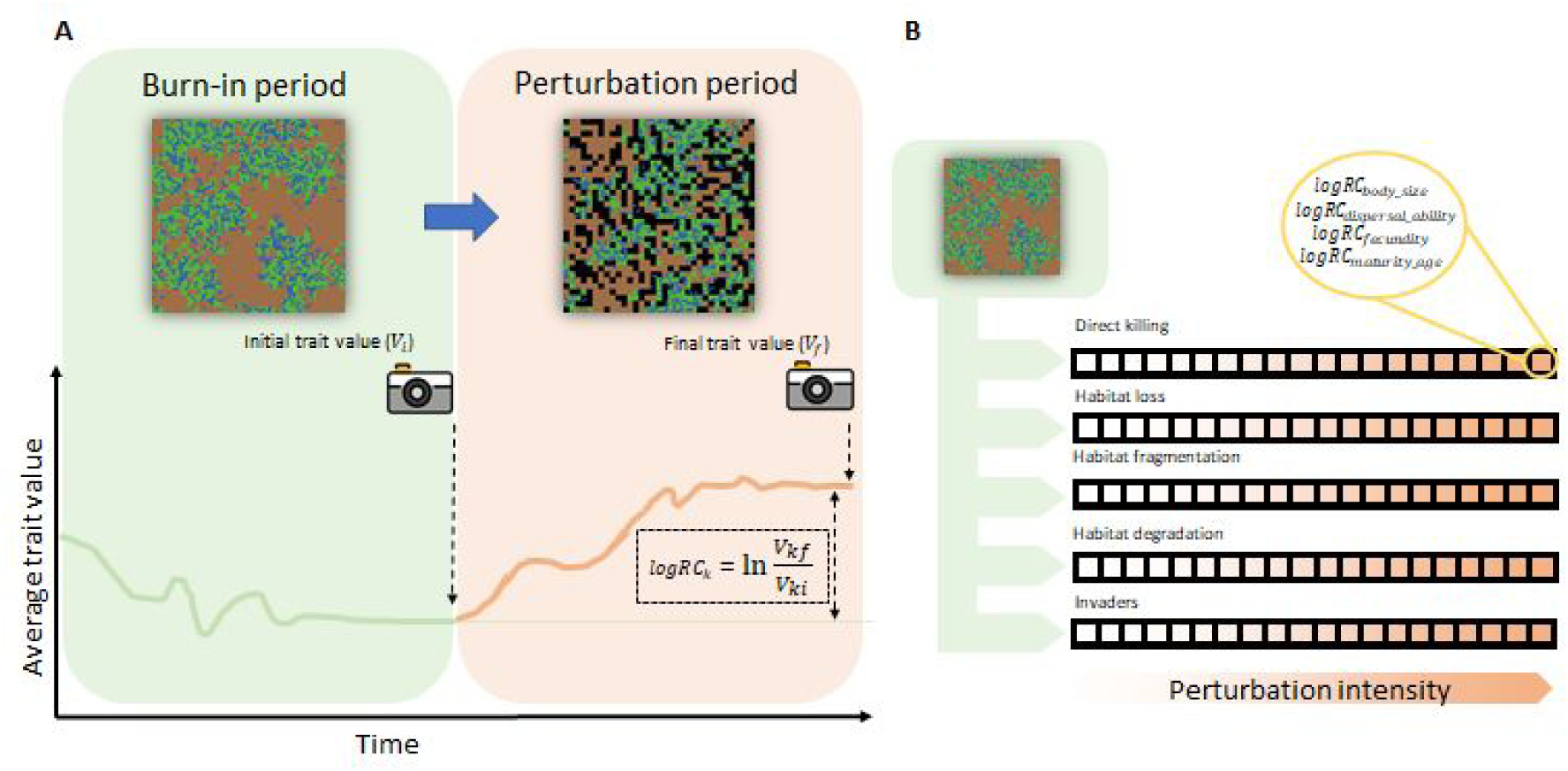
Schematic representation of the model design. **A)** Average trait value (any of body size, fecundity, maturity age, or dispersal ability) of a population of virtual organisms as a function of simulation time. As the model is initialized at time 0, natural selection and mutation in trait values shift the average trait values up and down, until a local optimum for each trait is attained. This period, in which the trait values are still varying over time, is called the burn-in period. Once the population reaches stable trait values, the burn-in period ends. At the end of the burn-in period, the model takes a snapshot of the mean trait values (initial trait values). The perturbation is activated, which may change the traits of the population of organisms. Once stable trait values are attained again, the simulation stops and another snapshot is taken. The relative change in trait values of each trait (logRC) is calculated as ln(initial value/final value). **B)** Experimental setting of the experiment. Each population is subjected to a single perturbation with a given intensity. Each perturbation at a given intensity was tested 64 times.

### The environment

The environment consisted of a continuous 2D space in which organisms foraged in the form of an underlying toroidal matrix of 33×33 patches (Fig. 2A). Patches were classified as either resource patches or bare patches. The resource patches generated resources linearly every timestep until they reached a maximum amount of resources. The quantity and spatial distribution of resource patches were controlled at the beginning of the simulation (Supplementary materials S1).

**Figure 2:**
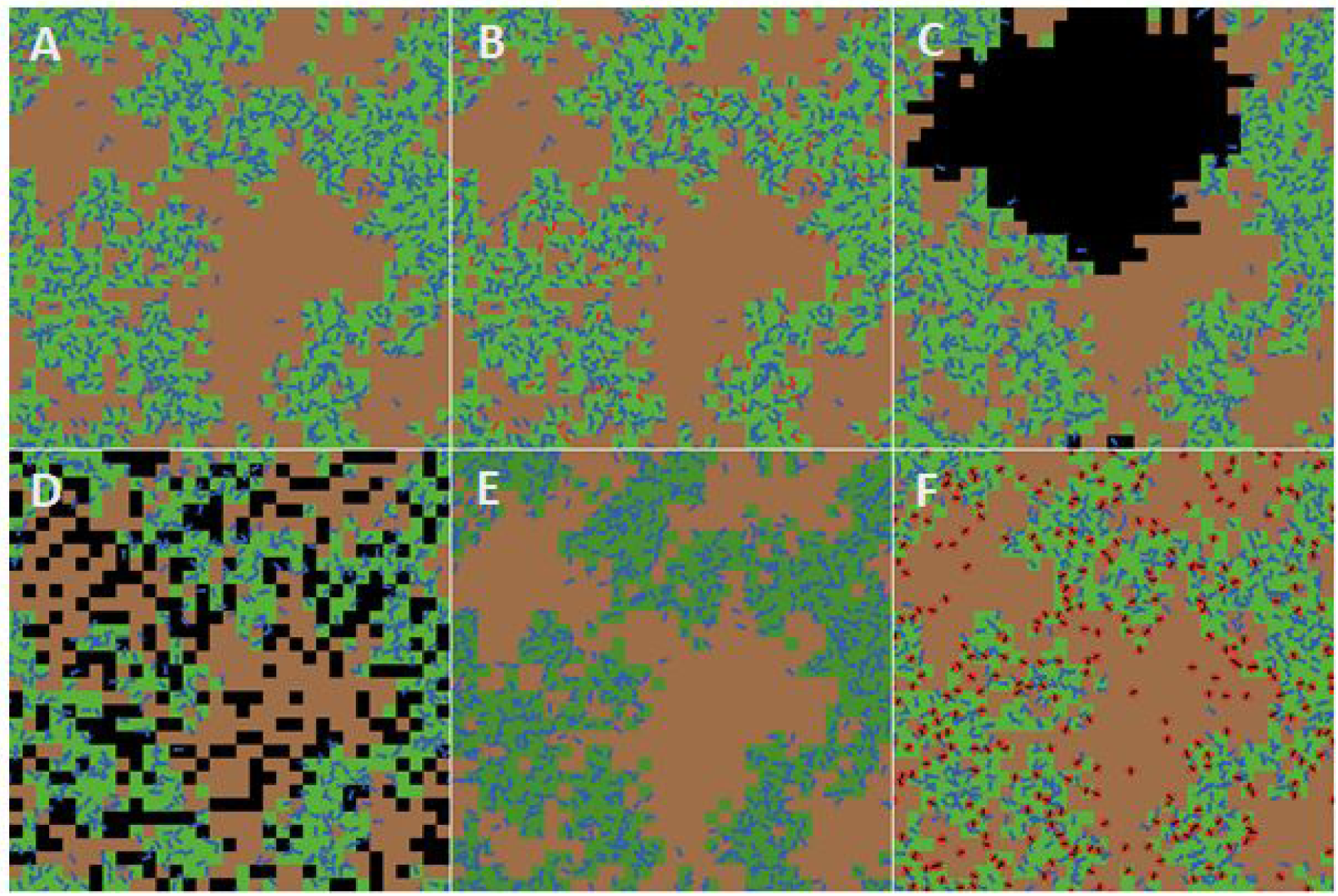
Visual representation of the same environment under different perturbation regimes. Blue “ bugs” are organisms, green patches contain resources, and brown patches contain no resources. A: No perturbation added. Organisms tend to cluster in regions where resources are available. B: Direct killing. Ten percent of organisms die at each round, marked in red. C: Habitat loss. A single fragment covering 30% of the cells was affected by habitat loss, without resources, marked in black. D: Habitat fragmentation. Thirty percent of patches were randomly assigned without resources, black patches. E: habitat degradation. All resource cells had a decrease in resource regeneration rate and in maximum resource value, indicated in dark green. F: Invaders. Invasive competing organisms were added, being immortal, not reproducing, but consuming resources, represented in red and black.

### The virtual organism

Organisms were mobile entities that reproduced asexually. They were characterized by an energetic currency, their age, their position in the continuous environment, and their focal traits. Organisms competed directly, accessing resources that fuel their maturation, reproduction, and survival needs (Fig. 3; Sibly *et al*. 2013; van der Vaart *et al*. 2016; Brown *et al*. 2018). Excess energy was stored in reserves up to a maximum determined by *body size*. An organism dispersed whenever the intake of resources at the current timestep was lower than its needs for maintenance and maturation/reproduction. If the amount of energy in the reserves reached a certain ratio of max-energy (*energy-to-reproduce*) and the organism was an adult, it reproduced, and part of its energy was allocated to its offspring.

**Figure 3:**
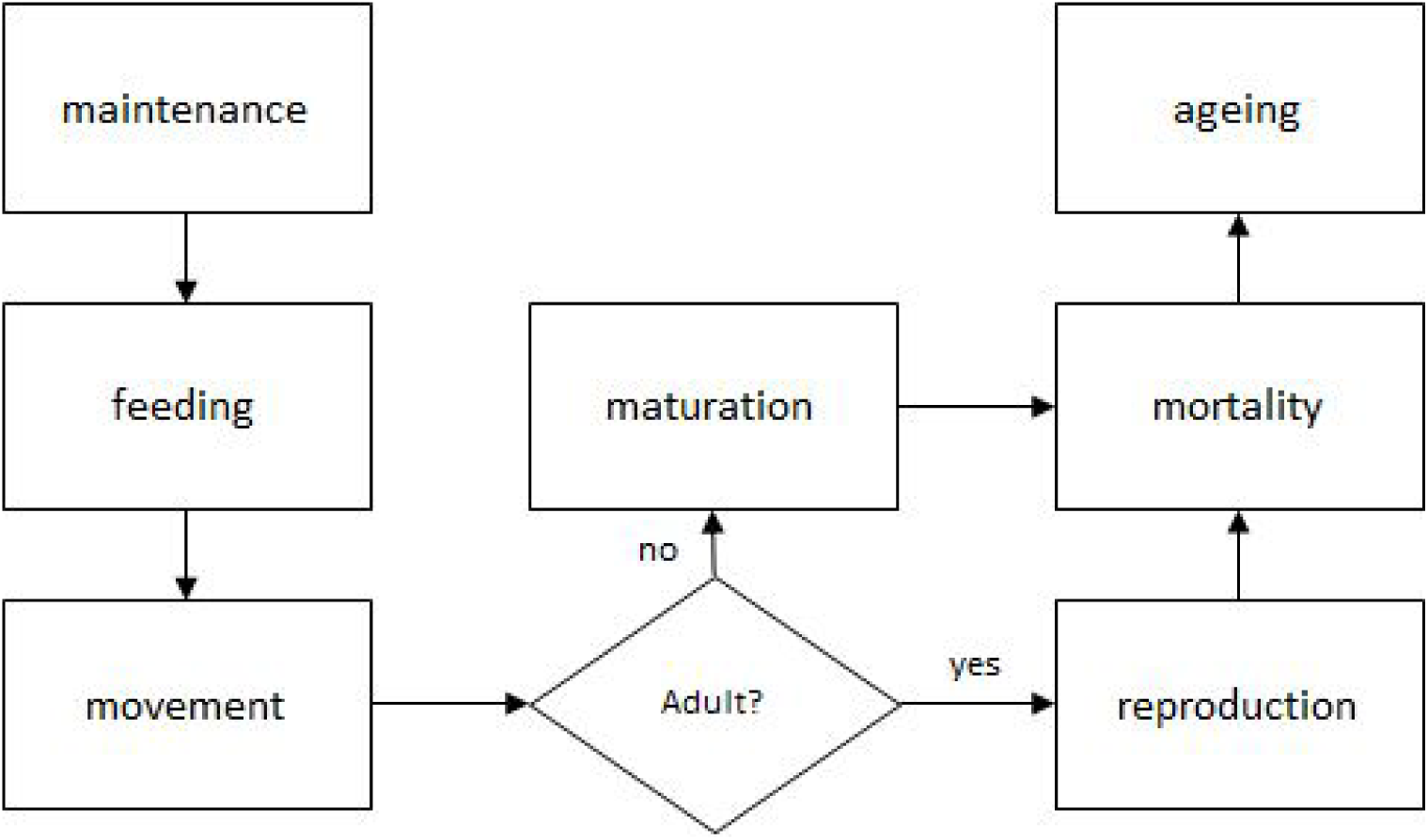
Functions executed by each organism at every time step. Each function’s details can be checked in the ODD protocol in Supplementary Materials S1.

Organisms died whenever the energy in reserves fell below 0, if they were killed by the perturbation *direct killing*, or due to old age. The focal traits influenced the fitness of organisms by determining the order in which they foraged (*body-size*), the volume of energetic intakes and expenditures of organisms (*body-size* and *dispersal-ability*), how quickly they reached maturity and died (*maturity-age*), how quickly, and thus how far, they could move in the environment (*dispersal-ability*), and how efficient they were in leaving progeny and spreading their traits in the population (*fecundity*).

#### Body size

Being larger than other organisms competing for resources is a competitive advantage for both animals and plants through access to more resources (Kingsolver & Pfennig 2004; Hone & Benton 2005; Kingsolver & Huey 2008). On the other hand, energetic expenditure and acquisition scale positively with the body size of organisms (Brown *et al*. 2004; Glazier 2005, 2010). Therefore, in this model organisms executed their functions in an energy-sorted order: organisms with more energy, which was positively correlated with body size, had higher chances of foraging first. Larger organisms intook and used energy at a faster rate than smaller organisms (Supplementary materials S1).

#### Maturity age

The ability to reach maturity earlier is accompanied by a reduced ability to correct mistakes when replicating DNA, which has multiplicative negative effects on cell functioning, limiting their ability to live longer (Kirkwood 1977; Dowling & Simmons 2009; Selman *et al*. 2012). In the model, longevity was proportional to the maturity age of an organism.

#### Fecundity

Investment in many offspring has a negative effect on each offspring’s survival shortly after birth (Fox & Czesak 2000). In the model, organisms with higher fecundity generated offspring with lower initial energy.

#### Dispersal ability

Higher dispersal ability may impose energetic, risk, and opportunity costs on organisms that may be expended during the pre-dispersal phase, transfer period, or resettlement phase (Bonte *et al*. 2012). On the other hand, organisms with higher dispersal ability can move farther away from their current position than poor dispersers. For simplicity, in the model the dispersal cost was debited for every dispersal event, and was proportional to dispersal distance (thus, there are energetic, and opportunity costs to be paid during transfer).Organisms in this model dispersed following a Brownian motion with random orientation, and with speed sampled from a uniform distribution with minimum zero and maximum *dispersal-ability*.

### Burn-in phase

A total of 500 organisms were added to the environment, initialized with random values of traits sampled from an exponential distribution. This ensured that a wide range of trait combinations was tested at the simulation start. As time advanced, the number of unique trait combinations started decaying, and due to mutation of trait values the simulations tended to stabilize around specific trait values. The model detected when a simulation stabilized by checking if there had been significant changes in trait values and the number of organisms over time (check ODD protocol in the Supplementary Materials S1 for further information about the stopping conditions of the algorithm). As the model triggered stable conditions, the burn-in phase ended, and the mean trait values of the population were recorded. Each of these stable populations was replicated and each replicate (64 total replicates) was subjected to different threat types and intensities (only one threat type and intensity per replicate).

### Threat phase

The threat phase started by subjecting a stable population to a threat. The threats in this model were built to mimic the main biodiversity extinction drivers (Table 1): direct killing, habitat loss, habitat fragmentation, habitat degradation, and invaders. The simulation ended when the mean trait values stabilized again (the stopping conditions test was called again). Each threat was applied following a gradient of intensity (only one per population, Fig. 1B). We set the maximum threat value as the value at which half of the populations per value went extinct before stabilizing in new trait values. We then created a gradient ranging from no threat (0) to the maximum value of each threat type in 20 steps. In total, we subjected 64 populations to each threat type and intensity.

**Table 1:** Types of threat studied, their description, and real-world equivalents.

| **Threat** | **Description** | **Examples** |
| --- | --- | --- |
| Direct killing | Removal of organisms from the environment | Fishing and hunting, introduced predators, lethal effects of pollution |
| Habitat loss | Destruction of habitat in a continuous area | Clearing of natural ecosystems for agriculture, urbanization, etc. |
| Habitat fragmentation | Destruction of habitat in a non-continuous area | Roads, agricultural landscapes with some remnant forests, edges of a forest being cleared |
| Habitat degradation | Deterioration of habitat quality | Landscape dominated by invasive plant species, with high nitrogen input from agriculture, affected by sources of pollution such as pesticides and light pollution |
| Invaders | Introduction of organisms with similar traits, competing for the same resources | Replacement of native species by functionally and phylogenetically similar introduced species. |

### Statistical analysis

The relative change in trait value (*logRC*_*k*_) was calculated using the following equation:

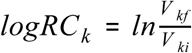

where *V* _*kf*_ is the mean value of the trait *k* in the population when the simulation ended, and *V* _*ki*_ the mean value of trait *k* of the initial population (before adding the threat). To calculate the signal of the relationship across all replicate populations and to see whether this signal was significantly different from zero (thus positive or negative), we performed linear mixed-effects models for each (threat, trait) pair. Since each replicate population was run at different threat intensities (see subsection “ Burn-in phase” above), we modelled the logRC within a linear mixed-effects model framework:

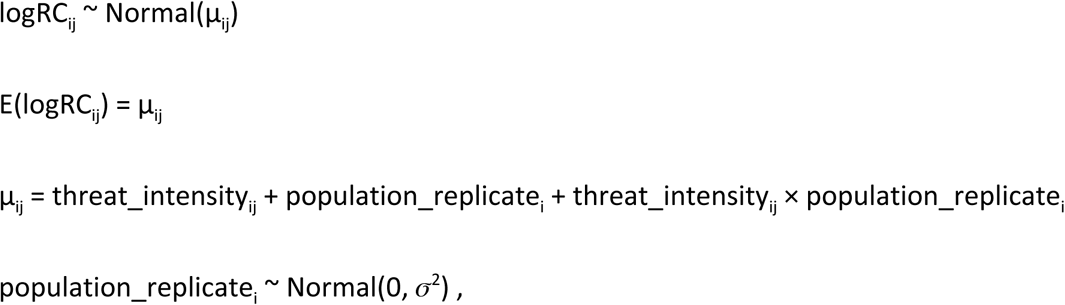

where logRC_ij_ is the *j*th observation of population replicate *i*, and population_replicate_i_ being the random intercept and slope, assumed to be normally distributed with mean 0 and variance σ^2^. We forced the intercept through the origin because under threat intensity = 0, logRC is expected to be 0. In those (perturbation, trait) pairs where the relationship was significant, we recorded the signal of the relationship.

As parameter values of the standard analyses (e.g. initial number of resource patches, energetic intake of organisms, and mutation rates of traits), we chose values that generated both realistic scenarios, and that allowed trait values to increase or decrease after adding threats (values presented in the Supplementary Materials S2). In addition to the standard analyses, we ran robustness tests (Supplementary Materials S2) to determine whether changing parameter values had a qualitative influence on the model outputs. We considered that the choice of values of a given parameter influenced the results if changing its value qualitatively modified the results, i.e., if we observed that the relationship between logRC and perturbation intensity was negative under a robustness experiment, while under the standard experiment values the relationship was positive, and vice-versa. Thus, we considered that qualitative changes occurred if there was a reversal in the sign of the relationship between the logRC and perturbation intensity. The scripts to run the Netlogo model and statistical analyses are available in Supplementary Materials S3.

## Results

Body size and maturity age decreased and fecundity and dispersal ability increased with increasing levels of direct killing of individuals (Fig. 4). Results were consistent across all robustness analyses for maturity age and dispersal ability (i.e. there were no robustness analyses results that led to opposite effects on traits, Table S3.1), but not for body size and fecundity (Supplementary Materials S2).

**Figure 4:**
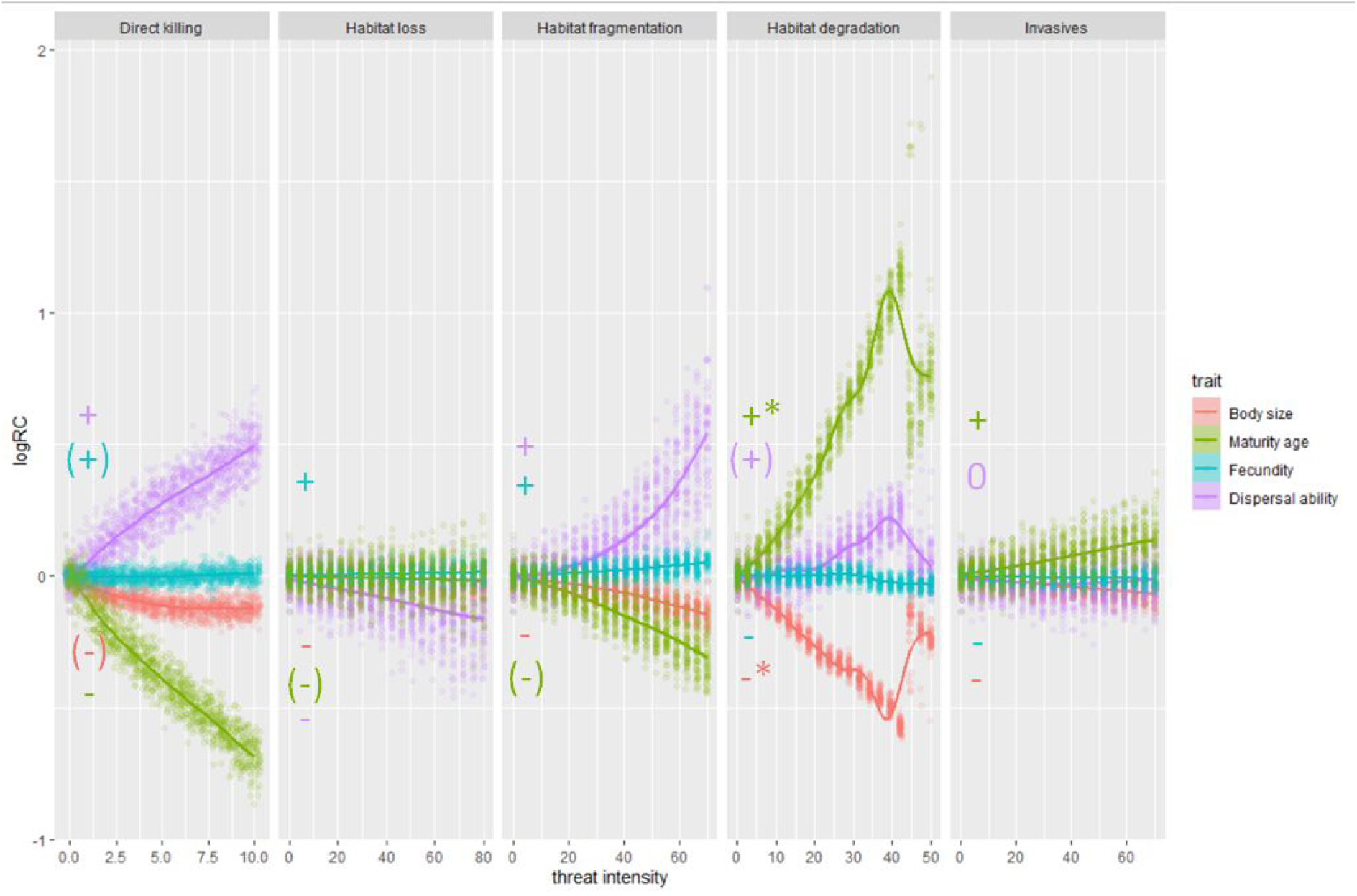
Evolution of the mean value of each trait as a function of perturbation intensity. The perturbation intensity was a gradient from zero to the value that allowed at least 50% of the simulations without species extinctions. Curves are “ loess” smoothers. Dots are individual replicates (total 64 replicates per treatment). “ +” and “ -” indicate slopes that are significantly larger and smaller than zero, “ ()” indicates results that were not robust to at least one parameter change (see Supplementary Materials S2), and “ *” indicates non-convergence of the linear mixed models, the signal being visually evaluated.

As a response to habitat loss intensity, body size, maturity age, and dispersal ability decreased, while fecundity increased. All changes were consistent independently of simulation parameters, except for maturity age, which showed a positive signal under a range of robustness tests (Table S2.1).

Habitat fragmentation led to mostly similar outputs as direct killing (Fig. 4). Body size and maturity age decreased, and dispersal ability and fecundity increased. All patterns were consistent independently of simulation parameters, except for maturity age, which under certain values showed a positive response (Supplementary Materials S2). Although this threat also implied habitat loss, the change in trait values was greater under habitat fragmentation, revealing the direct effects of fragmentation beyond those of loss.

Maturity age and dispersal ability increased with increasing levels of habitat degradation, although there was an inflection point at extreme levels of perturbation (which did not qualitatively have an impact on the direction of change). Body size and fecundity decreased, but again with inflection points in the intensity gradient (Fig. 4). All patterns were robust to parameter values, except dispersal ability, which decreased under certain scenarios (Table S2.1).

Adding competing invasive organisms with traits similar to the native led to increasing maturity age, and to decreasing body size and fecundity (Fig. 4). No change was detected in dispersal ability. All results were robust to parameter changes (Table S2.1).

## Discussion

Our work confirms in simulated settings that different threat types act differently on organisms depending on their traits. The way that organisms respond to threats or the probability with which species can go extinct may therefore depend on the specific threats to which they are subjected. Most patterns found were robust to different parameter values, meaning that the underlying mechanism should be consistent across many ecological systems in nature. We should note that, being a single-species experiment, our simulations did not test species extinction directly. They did test the mortality of individuals with different traits that make the species evolve, in this way indirectly testing extinction.

### Direct killing

Direct killing negatively affected large, slow-living, poorly fecund, and poorly dispersive organisms. Large sizes were removed from the environment because as organisms were killed at random, competition for resources diminished, which then reduced the need for the competitive advantage that larger size brings. Under some parameter value choices, however, the small organisms were instead disfavoured (Supplementary Materials S2). This seems to be due to the indirect effects of direct killing on resource availability. As organisms die, resources become more abundant. Abundant resources thus allow organisms to become larger under some parameter values.

Direct killing also heavily impacted organisms with slower life-cycles. Organisms taking longer to reach maturity faced greater probability of being killed before they were able to reproduce, and hence, the simulations became dominated by fast-living organisms.

Organisms with poor fecundity were disfavoured for most of the parameter values tested. As direct killing affected all organisms with the same probability, individual survival became less relevant than producing more offspring to compensate for the mortality. In simulations where organisms had a high longevity-to-maturity age ratio (due to low maturity age), they tended to evolve towards low fecundity, and hence, highly energetic offspring. These offspring quickly reproduce, reducing the chances of being killed before reproduction.

Our results also suggest that direct killing disfavours organisms with poor dispersal ability. This effect was again seemingly due to the increased resources available at each patch; as organisms died more frequently, more resources became available. These resources were better exploited by those organisms that found them first, or those with more dispersal ability.

The widely reported extinction of the megafauna before the 18th century has already shown that large-sized, slow-living, low fecund, and less mobile fauna are particularly susceptible, although it must be remarked that human hunting and fishing have long been selective towards large species (Diamond 1989). The same dual pattern seems to be occurring in recent times, at least for vertebrate taxa. Among fish and mammal species threatened by overexploitation, larger sizes tend to be more at risk (Olden *et al*. 2007; González-Suárez *et al*. 2013), a fact that is typically associated with their slow life-cycles. Additionally, threatened mammals facing overexploitation show lower fecundity independently of size effects than those not threatened (Strauss *et al*. 2006; Warzecha *et al*. 2016). Other sources of direct killing, such as the presence of an alien predator species (Matsuzaki *et al*. 2011) or disease (Murray & Hose 2005), could be affecting organisms with the same traits that we found, but thus far studies examining interactions between threats and traits have been limited (Murray *et al*. 2014). Interestingly, the increased vulnerability of small-sized individuals also emerged in a model evaluating the impact of predation on the size and maturity age of organisms, showing that prey size often increased when predation had an indirect effect on resource availability of prey (Abrams & Rowe 1996).

### Habitat loss

Our model suggests that habitat destruction has a negative impact on organisms with larger sizes, slow life-cycles, low fecundity, and high dispersal ability.

The decrease in body size could be explained by the reduction of total resource area in the environment. As habitat loss decreased the number of suitable habitat patches, it decreased the area of continuous suitable habitat, threatening larger size individuals that require abundant resources. Lower population sizes due to resource scarcity also incur greater probability of local stochastic accidents (Pimm *et al*. 1988). Edge effects further exacerbate this due to the high spatial resource unpredictability seen through the organism’s point of view.

Long-living organisms were also negatively affected by habitat loss. The higher spatial heterogeneity may negatively affect slow life-cycles, because these organisms attain reproductive age more slowly, and thus, reproduce less often, not compensating for mortality losses due to movement into the matrix of the unsuitable habitat. Under some robustness scenarios, though, habitat loss was detrimental to fast life-cycles. Typically, slow-lived populations survive local extinction better at lower population densities than fast-lived populations (Pimm *et al*. 1988), and with less stochastic fluctuations, thus possibly conferring higher resilience in spatially heterogeneous landscapes.

Habitat loss intensity was detrimental to organisms with lower fecundity. High fecundity of organisms in the model came at the cost of decreased survival of individual offspring. When faced with a reduction of habitat, however, individual survival is offset by increased edge effects and spatial unpredictability.

Organisms with larger dispersal ability were negatively affected by habitat loss. As habitat decreases in size without alternatives to which to move, individuals that invest in long-distance dispersal incur higher mortality.

The impacts of habitat loss on species’ traits have not been studied extensively. Conceptually, larger sizes are thought to be the most susceptible to decreases in fragment area (Ewers & Didham 2006). However, empirical evidence supporting the higher risk faced by larger organisms is very limited, due to the existence of many confounding factors under the umbrella of body size (Ewers & Didham 2006); large size may correlate with other traits that increase (e.g. dispersal ability (Warzecha *et al*. 2016)) or decrease (e.g. trophic level (Henle *et al*. 2004)) the organism’s capability of surviving in areas with increased habitat loss. Likewise, the effects of habitat loss on lifespan have not been conclusive. Plants with greater longevity were hypothesized to be better at coping with habitat isolation (Lindborg 2007), as they face less fluctuations in their population levels, but fast life-cycles have been shown to survive well in isolated fragments, likely because fast life-cycles were also characterized by being very fecund (Lindborg *et al*. 2012; Marini *et al*. 2012). In a grassland that has faced extensive habitat loss, surviving plants in isolated habitat patches had greater seed output than the original species population prior to the habitat loss event (Saar *et al*. 2012), which authors associated with their probable better ability to find microsites than species with lower seed output. A meta-analysis indicates that butterflies with poorer fecundity are more affected by habitat fragmentation (Öckinger *et al*. 2010). As for dispersal ability, evidence shows that when patches of remaining habitat are very isolated, organisms with reduced dispersal ability often cope better (Henle *et al*. 2004; Ewers & Didham 2006; Saar *et al*. 2012). However, isolation and no dispersal may have consequences for genetic drift and erosion for organisms reproducing sexually (Zambrano *et al*. 2019) and for species’ ability to respond and shift their geographical ranges in the face of climate change (Hodgson *et al*. 2011).

### Habitat fragmentation

Like habitat loss, habitat fragmentation had a negative effect on large, slow-living, and poorly fecund organisms, but a negative effect on poor dispersers. At the same proportion of habitat removal, the effect of habitat fragmentation on traits was stronger than that of habitat loss, revealing the importance of these two effects combined to drive population declines. Slow-moving organisms, being favoured under the pure habitat loss scenario were disadvantaged under habitat fragmentation, as they could not move among a complex matrix of suitable and unsuitable patches.

With increased fragmentation, organisms with higher dispersal that could move to new, relatively easy-to-find patches, were likely to benefit, also avoiding the dangers of low population size and stochasticity in very small fragments.

Increasing fragmentation has been shown to benefit good dispersers. For example, species of butterflies and moths in fragmented landscapes were characterized by greater dispersal ability (Öckinger *et al*. 2010). Differences in dispersal ability among landscapes can be found even within species, whereas isolated patches in fragmented landscapes were more frequently visited by larger, more mobile wild bees (Steffan-Dewenter & Tscharntke 1999; Warzecha *et al*. 2016).

Studying the impact that habitat loss and fragmentation have on species traits is challenging. First, because often traits correlate with each other. Second, because the literature on the effects of habitat loss and habitat fragmentation on species’ traits is fuzzy, since these terms are often used interchangeably (Tscharntke *et al*. 2012; Fahrig 2017; Zambrano *et al*. 2019). Third, because habitat loss and fragmentation can be subdivided into five components, each of which is hypothesized to have different effects on species’ life-histories. These components comprise a decrease in fragment area, distance from the edge, shape complexity, isolation of the remaining habitat, and contrast between suitable and non-suitable habitats (Ewers & Didham 2006). For example, the decrease in fragment area is thought to negatively affect mostly the intermediate dispersers, whereas the long-distance dispersers can find suitable patches elsewhere and poorly dispersers do not incur unnecessary dispersal events that lead to increased mortality (Henle *et al*. 2004; Ewers & Didham 2006).

### Habitat degradation

Habitat degradation in our model largely equates to resource scarcity. Larger organisms require more resources, therefore being especially vulnerable to reductions in resource quantity. In the model, organisms that mature more quickly survive less time as adults. Under these settings, the chances that they may be able to accumulate enough energy to reproduce are smaller. Therefore, greater longevity is favoured, as it gives a chance to accumulate the energy required for reproduction. In the same way, by concentrating more energy onto only a few offspring, organisms increase the odds that their offspring will survive famine during the juvenile stage. Organisms that are poorer dispersers are disfavoured. As resources become less abundant, the necessity to rapidly find these elsewhere increases.

The effect of habitat quality as related to resource availability on biodiversity variables at a landscape level has been poorly studied, despite the fact that along with habitat loss and fragmentation, it is one of the major drivers of species extinction (Mortelliti *et al*. 2010).

Furthermore, isolating the effect of habitat degradation *per se* from other landscape-level drivers of change is difficult, as it often comes as a side-effect of habitat loss and fragmentation (Fischer & Lindenmayer 2007; Mortelliti *et al*. 2010). Temperate ungulates are examples of animals that have developed slower life-cycles in response to lower resource density, while minimizing their reproduction (Skogland 1985; Ferguson 2002). For instance, when food is scarce, pregnant wild deer abandon their fetuses in preference for their own survival (Skogland 1985). Plants living in nutrient-poor scenarios, usually have traits that confer low growth rates, such as high root-to-stem ratios and low specific leaf area (Grime 1977). Likewise, cave organisms survive extreme resource-depleted scenarios. They are invariably long-lived, with low fecundity, and large organisms simply cannot survive entirely in caves due to lack of resources. All of these traits were favoured by our simulations, which replicate similar circumstances to those frequently occurring in degraded habitats. Our understanding of the mechanisms through which habitat degradation affects species is far from clear since, as we do not understand the types of traits that are selected (Mortelliti *et al*. 2010), even though habitat quality may typically outweigh the importance of habitat loss and fragmentation to species extinctions, as suggested by other agent-based simulations (Heinrichs *et al*. 2016).

### Invaders

The effect of invasives was somewhat similar to habitat degradation: a reduction in body size and fecundity, and an increase in maturity age (but no effect on dispersal ability).

The invasive organisms added competition for the same resources, reducing the number of resources available and causing similar outcomes for most traits. Organisms become smaller as a direct response to decreasing resource availability, compromising their ability to access resources first due to their smaller size than invasive organisms. To compensate for lower resource intake, they become longer-lived so as to accumulate resources over a longer period of time before reproducing.

Additionally, they maximize their offspring’s survival by putting more energy into a few, rather than many, offspring. Invasive species had, however, no effect on the dispersal ability of natives. Under habitat degradation, the population density of organisms is reduced.

Therefore, higher dispersal may be useful to find other patches of suitable habitat. Such is no longer the case with the addition of an invasive species. The density of native and invasive organisms combined does not change much in relation to the pre-invasive event, and therefore, if an organism disperses to another patch it may find exactly the same density of organisms, curtailing the advantage of moving fast.

The impact of invasive competitors on the traits of native species competing for the same resources has not been examined thoroughly. A comparative study of extinction risk compared the traits of freshwater invasive, threatened native, and non-threatened native fish species, and found that threatened natives had significantly faster life-cycles and higher fecundity than those of non-threatened native species (Liu *et al*. 2017). The authors suggested that the reduced fecundity and faster lives could be related to higher environmental stochasticity, and hence reduced resilience in the face of environmental change. River systems are especially threatened by the presence of invasive species; in many regions the number of invasive fish species surpasses a quarter of the total species richness (Leprieur *et al*. 2008), and in this system in particular they could be the number one cause of extinctions (Light & Marchetti 2007). Therefore, the faster life cycle of threatened fish species could be a consequence of the presence of alien competitors. A meta-analysis of trait differences between invasive and non-invasive plant species has shown that natives are on average smaller and have lower growth rates than invaders (Kleunen *et al*. 2010). Trait values of successful native plant species were however similar to successful invaders in another study of invasiveness across many habitats in Central Europe (Loiola *et al*. 2018), and reflect the fast growth capabilities needed to deal with increased disturbance in their environment (Leishman *et al*. 2010).

## Conclusions

In this study, we explored how widely the examined drivers of extinction can differentially affect individuals (and species) with contrasting traits. Notably, we have shown that different threats have different outcomes, even in such simple scenarios as the ones we tested. For all the scenarios tested, threats that directly induce increased mortality to populations are detrimental to species with slow life-cycles and poor dispersal ability.

Threats that decrease the amount of habitat are harmful to larger organisms with poor fecundity. The effect of dispersal ability mostly depends on the spatial configuration of the threat. Threats that reduce the quality and quantity of resources in a landscape are harmful to species with a reduced capacity to compete under low resource conditions such as those that are large sized, and have rapid life-cycles and high fecundity.

Most patterns found in the simulations are congruent with previous literature using empirical data across varied taxa and regions. Other patterns found were not previously reported, either due to lack of data or, more often, due to confounding factors that do not allow independent tests of threats and traits. For example, discriminating the effects of habitat fragmentation from those of habitat loss alone is difficult. Nevertheless, it is important to separate them, as these two drivers have opposite effects on the dispersal ability of organisms. Additionally, the effect of different threats on body size is typically difficult to assess because this trait is often correlated with many other traits. We provide a baseline hypothesis for body size, that is; large size is detrimental to species for any given threat, independently of other traits, due to the high resource needs that large sizes incur to species. These examples suggest future avenues for research and testable predictions that can be confirmed or disproved as proper data become available.

## Acknowledgements

We thank Volker Grimm for useful discussion and technical support during a previous version of the model. F.C. and P.C. were funded by Kone Foundation, Finland, with the project ‘Trait-based prediction of extinction risk’. L.C. was partially supported by UIDB/04046/2020 and UIDP/04046/2020 Centre grants from FCT, Portugal (to BioISI).

## Supplementary Materials S1: ODD Protocol

### 1. Purpose and patterns

The model is designed for theoretical exploration of biological trait value changes in response to five perturbation types: direct killing, habitat loss, habitat fragmentation, habitat degradation, and invasives. Specifically, we will explore changes in the values of four traits (body size, maturity age, fecundity, and dispersal ability) of virtual organisms moving and competing in virtual landscapes. From the resulting trait changes implications for extinction risk will be discussed. The generic patterns used to claim that the model is internally consistent and realistic enough for its purpose are previously published trends in trait-extinction risk relationships.

### 2. Entities, state variables, and scales

The model’s entities are organisms, patches, and the environment. Their state variables are listed in Table S1.1. The model parameters are listed in Table S1.3. Organisms are mobile entities that move, consume resources, have a basic energy budget, reproduce asexually, and die. All organisms are of the same species. Variability in trait values is created through mutation and at start, in which organisms are assigned with random trait values. Patches are square units of space which can grow resources or not. The environment is constant during a burn-in period while average trait values of the population stabilize. The environment is then subjected to one of five different perturbation types: direct killing, habitat loss, habitat fragmentation, habitat degradation, or invasive organisms.

**Table S1.1:** Variables, their names, types, and units.

| Netlogo name | Description | type, unit |
| --- | --- | --- |

| Organism |  |  |
| --- | --- | --- |
| *Dynamic state variables* |  |  |
| energy | current energy | real, arbitrary unit of energy |
| age | age | discrete, timestep |
| energy-intake | Energy income of current timestep | real, energy |
| heading | Orientation | integer, degrees |
| xcor, ycor | Location | continuous, distance from center of environment |
| age-at-reprod | Age at which the last reproductive event happened | discrete, timestep |
| *Static state variables (taken from probability distributions)* |  |  |
| body-size | Body size | real, arbitrary unit of mass |
| maturity-age | age at which organism reaches reproductive potential | integer, timestep |
| fecundity | number of offspring during<br>reproductive event | integer, offspring |
| disp-ability | maximum distance traveled in<br>a timestep | real, patch length |
| *Derived variables (not state variables because they are calculated from other variables and parameters)* |  |  |
| max-energy | body-size *<br>metabolic-allometric-exponent | real, energy |
| max-energy-intake | max-energy-intake-coef *<br>max-energy | real, energy |
| maintenance-cost | maintenance-cost-coef *<br>max-energy | real, energy |
| maturation-cost | maturation-cost-coef *<br>max-energy | real, energy |
| dispersal-cost | dispersal-cost-coef *<br>max-energy | real, energy |
| energy-to-reproduce | energy-to-reproduce-coef *<br>max-energy | real, energy |
| energy-after-reprod | energy-after-reprod-coef *<br>max-energy | real, energy |
| longevity | maturity-age *<br>longevity-maturity-coef | discrete, timestep |
| **Patches** |  |  |
| *Dynamic* |  |  |
| max-resources | Maximum resources | real, energy |
| resources | Energy/resources | real, energy |
| regen-rate | Rate of energy/resource<br>regeneration | real, energy |
| **Environment (stopping conditions)** |  |  |
| *Dynamic* |  |  |
| next-checkpoint | Number of timesteps until the next check of stopping conditions | discrete, timestep |

We chose to represent growth in terms or age, which corresponds with potential energy accumulation, but left body size, in the sense of the final, achievable body size, as a constant trait.

One timestep has no real unit of time equivalent, but can be delineated by considering a “ minimum resolution” which is the necessary amount of time for an organism to perform any of these functions: feeding, maturation (if juvenile) or reproduction (if adult), and movement. The total extent of time of a simulation is very variable, because it depends on how quickly the mean trait values of the population converge before, and after a perturbation. Convergence of trait values can be attained after only 500 timesteps, but typically simulations last between 1000-10000 timesteps.

The model world consists of 33⨯33 patches, which is large enough to generate maps with realistic spatial autocorrelation. We confirmed the robustness of our finding with both smaller and larger world sizes (Supplementary Materials S2).

### 3. Process overview and scheduling

Every timestep each organism executes the following submodels that are explained in detail in ODD section 7, “ Submodels” : **<u>maintenance</u>**, in which it deducts energetic costs of maintaining its body, and die if its energetic level becomes lower than zero; **<u>feeding</u>**, in which it obtains *energy* from the patch where it is; **<u>movement</u>**, in which it moves if the *energy intake* of the current timestep is lower than the predicted costs of maturation and maintenance; if they have not yet reached maturity age, **<u>maturation</u>** is executed, in which it spends energy in maturation; if maturity age was reached and enough energy is available, **<u>reproduction</u>**, in which the organism reproduces asexually; then **<u>mortality</u>** is executed, in which the organism may die due to the direct killing perturbation (during the perturbation phase), or due to old age; and then age (**<u>ageing</u>**). Each time step, the order by which the organisms execute their sequence is ordered according to the organisms’ current energy level, with some noise (**<u>sort-organisms</u>**), this way favoring organisms with higher energy levels on average.

Next, *resources* on patches re-grow (**<u>update-patches</u>**). At the end of each time step, in **<u>check-stopping-conditions</u>** it is checked if the number of organisms and the mean values of organisms’ *body size, fecundity, maturity age*, and *dispersal ability* have not changed considerably in relation to previous time steps. The first time that the stopping conditions are triggered the burn-in period ends, the mean trait values recorded, and the current state of the simulation is saved if <u>save-world?</u> is true. The threat phase then begins with **<u>add-perturbations</u>**, in which one perturbation is activated. If the stopping conditions are met a second time the whole simulation ends and mean trait values are recorded again.

When an offspring is born, its *body size, fecundity, maturity age*, and *dispersal ability* are inherited from the parent, but change due to mutation. Under **<u>organism-initialization</u>** high level variables are calculated, such as energy budgets considering the agent’s body size and system level energetic parameters and the longevity dependent on maturity age.

### 4. Design concepts

The sections Learning, Objectives, Prediction and Collectives do not apply to this model.

#### Basic principles

Four core assumptions govern the trade-offs between the focal traits (Table S1.2).

**Table S1.2:**
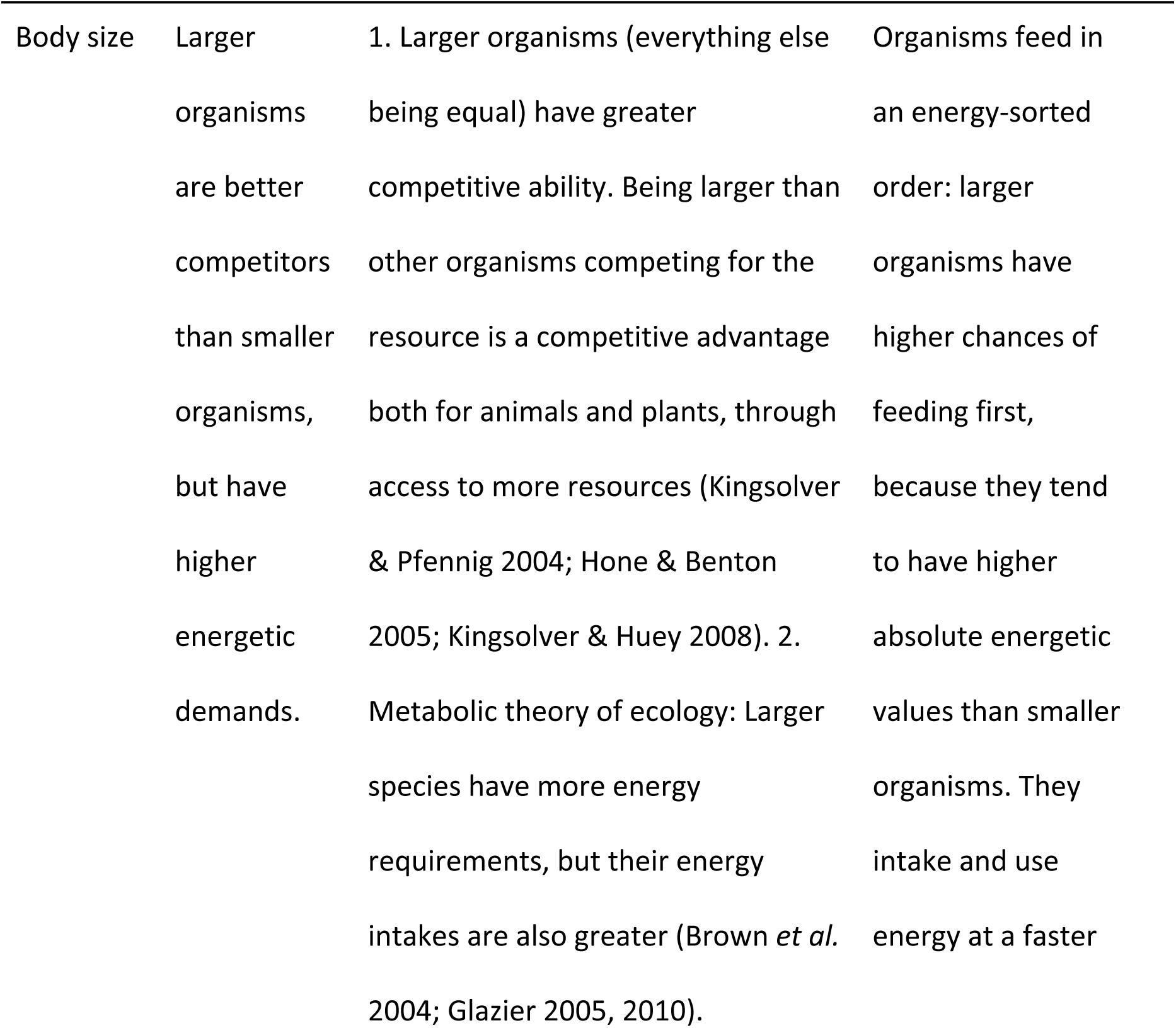
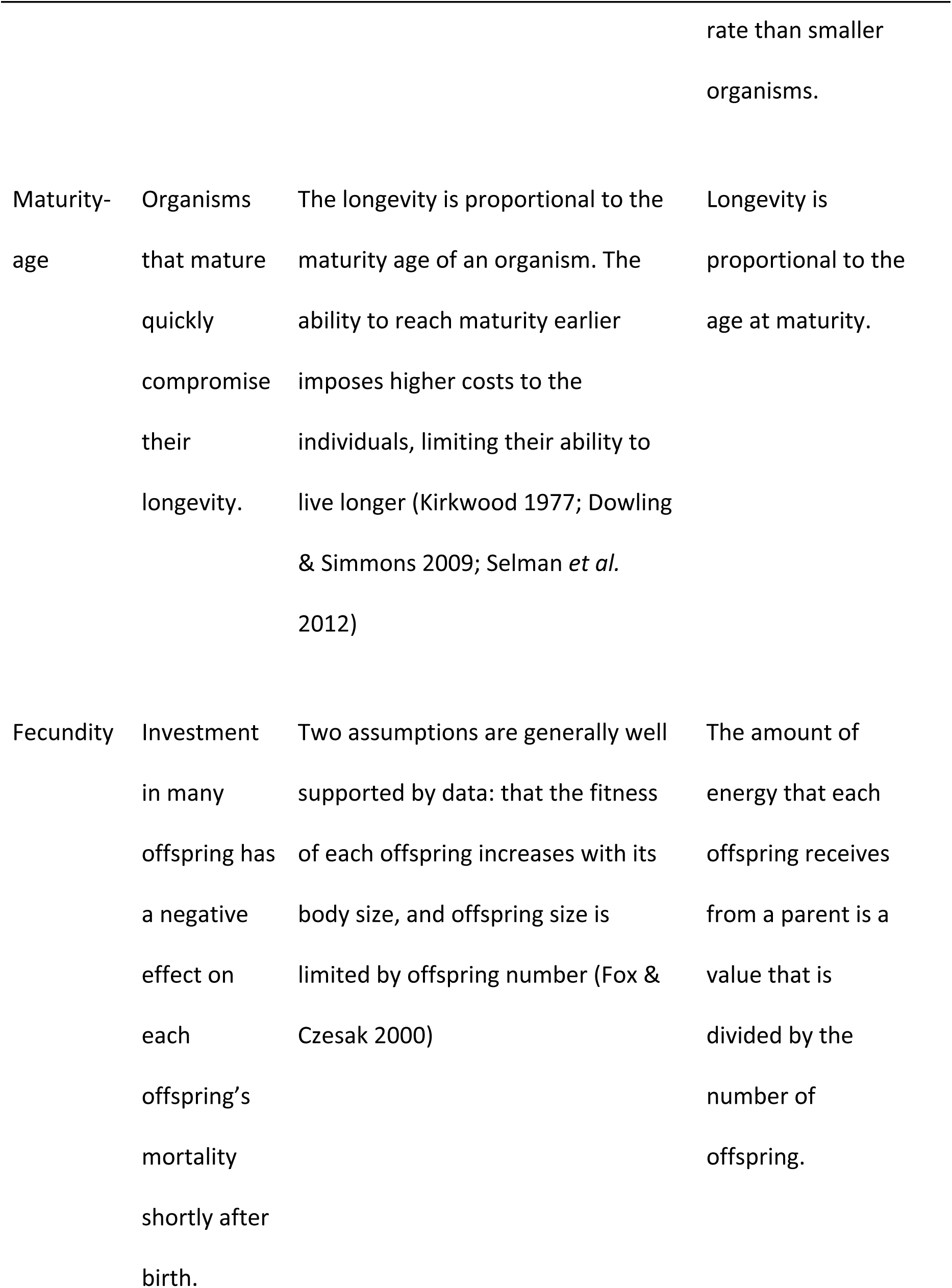
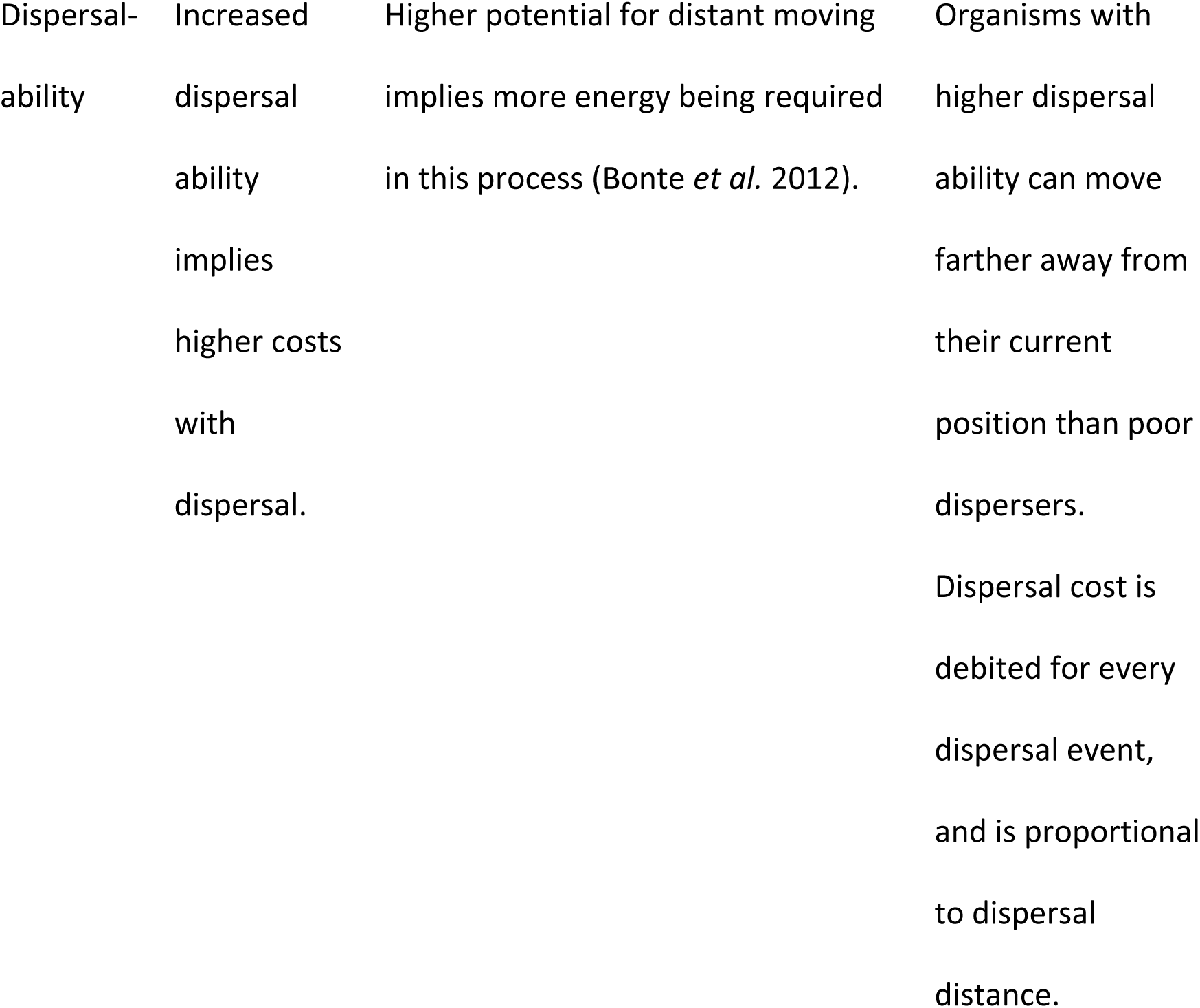
Main assumptions and theories underlying trade-offs in focal traits.

| Trait | Assumptions | Basis for the assumption | Implementation |
| --- | --- | --- | --- |
| Body size | Larger organisms are better competitors than smaller organisms, but have higher energetic demands. | 1. Larger organisms (everything else being equal) have greater competitive ability. Being larger than other organisms competing for the resource is a competitive advantage both for animals and plants, through access to more resources (Kingsolver & Pfennig 2004; Hone & Benton 2005; Kingsolver & Huey 2008). 2. Metabolic theory of ecology: Larger species have more energy requirements, but their energy intakes are also greater (Brown *et al.* 2004; Glazier 2005, 2010). | Organisms feed in an energy-sorted order: larger organisms have higher chances of feeding first, because they tend to have higher absolute energetic values than smaller organisms. They intake and use energy at a faster rate than smaller organisms. |
| Maturity-age | Organisms that mature quickly compromise their longevity. | The longevity is proportional to the maturity age of an organism. The ability to reach maturity earlier imposes higher costs to the individuals, limiting their ability to live longer (Kirkwood 1977; Dowling & Simmons 2009; Selman *et al.* 2012) | Longevity is proportional to the age at maturity. |
| Fecundity | Investment in many offspring has a negative effect on each offspring's mortality shortly after birth. | Two assumptions are generally well supported by data: that the fitness of each offspring increases with its body size, and offspring size is limited by offspring number (Fox & Czesak 2000) | The amount of energy that each offspring receives from a parent is a value that is divided by the number of offspring. |
| Dispersal-ability | Increased<br>dispersal<br>ability<br>implies<br>higher costs<br>with<br>dispersal. | Higher potential for distant moving<br>implies more energy being required<br>in this process (Bonte *et al.* 2012). | Organisms with<br>higher dispersal<br>ability can move<br>farther away from<br>their current<br>position than poor<br>dispersers.<br>Dispersal cost is<br>debited for every<br>dispersal event,<br>and is proportional<br>to dispersal<br>distance. |

#### Emergence

Our model is designed to let trait distributions and combinations emerge in response to the environment and changes therein. Organisms have different sets of traits from each other due to mutation during reproduction. During the burn-in phase natural selection excludes combinations of trait values that are not successful, due to the higher reproductive output of those that were successful. Over time, the population converges to an equilibrium set of trait values, which is a local optimum. When a threat is introduced, new combinations of trait values may become more successful than the ones at the end of the burn-in stage and therefore the trait values of the population shift towards a new equilibrium. The successful set of traits depends highly on the availability of resources and spatial arrangement of resources in the world, as well as on the intensity and type of the perturbation.

#### Adaptation and Sensing

Organisms perceive their internal energy/reserves level and energy-income and decide to move if their energy-income was lower than the costs of maintenance and growth.

#### Interaction

Interactions between individuals are indirect, via competition for a limited resource.

#### Stochasticity

Stochasticity is used:

- when generating a new resource distribution
- when mutating traits
- when dispersing (orientation and distance of organisms is random)
- sorting organisms at the beginning of each new time step
- selecting which patches are affected by the perturbations habitat loss, habitat fragmentation and habitat degradation
- selecting which organisms or patches are affected by background mortality
- selecting the trait values of invasive organisms.

#### Observation

Mean and standard deviation of all focal traits are collected at the moment before adding the perturbations and when the simulation ends. Furthermore, when testing specific modules, the values of all state variables may be monitored at every timestep.

### 5. Initialization

Initialization begins with generating a map that determines the spatial distribution of resources in the map (**generate-map**), and the values of state variables of patches with resources. All patches with resources are arbitrarily initialized with *max-resources* = 1, and resource-regen = 0.2. The absolute values of *max-resources* and *resource-regen* are not relevant, but the relative values between them determine the time in timesteps that it takes for a patch to go from 0 *resources* to *max-resources*. Higher values of *resource-regen* relative to the *max-resources* means that resource patches reach their maximum level of resources faster.

To allow a large range of different trait combinations for the initial trait values, and to avoid the mean trait values to converge very quickly towards a local optimum, each organism’s trait values are sampled from an exponentially distributed random floating-point number using Netlogo’s native procedure *random-exponential* with mean parameter 2. The number of initial organisms is arbitrarily set to 500.

Alternatively, the model can be initialized already from the beginning of the perturbation phase, in which a previously saved “ burned-in” model is loaded, if <u>load-world?</u> is true. <u>filename</u> identifies the character string under which the model was saved on disk. If <u>load-world?</u> is false, a new model begins anew.

### 6. Input data

This model uses no external data.

### 7. Submodels

To ensure that each submodel was working properly we made use of the Netlogo features such as inspection of agent’s values of state and other variables, the use of monitors to see the evolution of the trait values, and other useful information, such as the number of organisms, total amount of resources available, mean age at reproduction, etc, as well as the visualization tool incorporated, that allows the user to inspect each agent’s action. Under each of the submodels we provide some examples of problems we faced while implementing some of the original code.

#### generate-map

This submodel creates a landscape with dimensions given by the parameters <u>w-width</u> and <u>w-length</u>, and distributes resources to patches according to the parameters <u>nr-resource-clusters</u> and <u>resource-patch-fraction</u>. An initial number of patches, given by <u>nr-resource-clusters</u>, is assigned as resource patches, their <u>max-resources</u> assigned to 1, and their <u>resource-regen</u> initialized as <u>regen-rate</u>. The patches around each resource-patch are then also assigned as resource patches, until the number of resource patches in the environment reaches the total number of patches * <u>resource-patch-fraction</u>. The remaining non-assigned patches are then initialized as bare-patches, with <u>max-resources</u>, <u>resource-regen</u>, and <u>regen-rate</u> as 0. The parameter <u>map-seed</u> allows the creation of landscapes that are replicable by setting its value to a number larger than 0.

##### Rationale

*As <u>resource-patch-fraction</u> increases, the number of patches with resources increases. The more <u>nr-resource-clusters</u>, the less spatial aggregation between resource patches. If nr-of-resource-clusters is 1 resource patches will tend to aggregate in a single area, whereas when <u>nr-resource-clusters</u> is very high their spatial distribution is completely random and non-correlated (Fig. S1.1). Similar algorithms are used throughout the literature to simulate landscapes with different degrees of patchiness of resources (Fahrig 1998; Milles et al. 2020)*.

**Fig S1.1:**
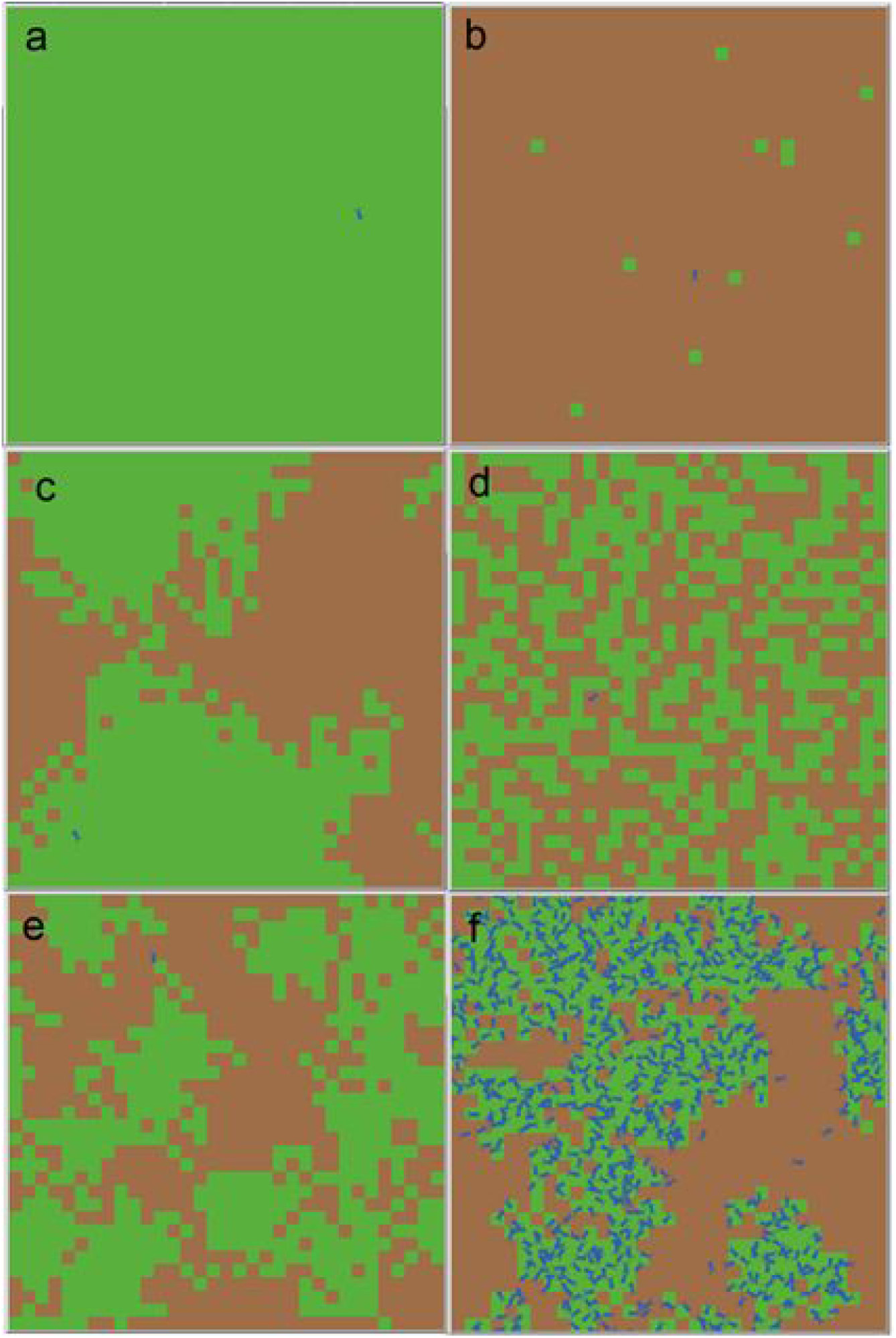
Example of maps generated by the submodel generate-map: the extremes of *resource-patch-fraction*, a) 1 and b) 0.01; the extremes of spatial aggregation, with *nr-resource-clusters* c) 1 and d) 544; the default values of *resource-patch-fraction* and *nr-resource-clusters* of 0.5 and 20 before the simulation starts e) and when the average values of traits in the population converges f). Green patches indicate patches with resources, and brown patches without resources.

#### organism-initialization

When organisms are born or initialized when simulation begins, derived variables of organisms are calculated (Table S1.1).

*Rationale: Derived variables are not strictly necessary in the model, in the sense that they are just the product of other variables and parameters. But their calculation once in a simulation makes the code faster*.

The maximum amount of energy of organisms (<u>max-energy</u>) is initialized as <u>body-size</u> ^ <u>metabolic-allometric-exponent</u>. Energetic balances, representing the maximum intake of energy during a timestep (<u>max-energy-intake</u>), maintenance and maturation costs per timestep (<u>maintenance-cost</u>, <u>maturation-cost</u>), and costs of dispersal per unit of distance traveled (<u>dispersal-cost</u>) are calculated in relation to their <u>max-energy</u>: <u>max-energy-intake</u> = <u>max-energy</u> * <u>max-energy-intake-coef</u>, and so forth (Table S1.1). Thresholds of reproduction, such as the minimum energy to reproduce (<u>energy-to-reproduce</u>), and the energy remaining in the organism after reproducing (<u>energy-after-reprod</u>), are calculated in the same way, by multiplying <u>energy-to-reproduce-coef</u> and <u>energy-after-reprod-coef</u> to <u>maximum-energy</u>.

*Rationale: Energy expenditures and intakes of organisms are directly linked to their body size. They are proportional to body size to the power of an allometric constant, typically 0.75, but varying between 0.66 and 1 (Brown et al. 2004). As such, in this model both the total energetic pool of organisms (<u>max-energy</u>), their expenditures and intakes of energy, as well as the amount of energy varies allometrically with body size*.

The longevity of organisms is also calculated as the product of <u>longevity-maturity-coef</u> with <u>maturity-age</u>.

#### update-patches

Patches regrow <u>resources</u>. <u>Resources</u> are incremented by <u>resource-regen</u> * <u>max-resources</u>, but not exceeding <u>max-resources</u> (in this case <u>resources</u> become <u>max-resources</u>).

*Rationale: In the real world, resources typically regenerate until reaching a maximum level. The rate at which resources regenerate may depend greatly on the system being studied. When resources are organisms, such as plants, their growth is typically modelled with logistic growth curves. Some systems are modelled with adapted logistic growth curves, such as vegetation under predation effects from herbivores. In such a case, the existence of below-ground reserves enhances growth at low aboveground biomass* (Mortensen *et al*. 2018). *Linear growth of resources may be modelled for systems with constant input of nutrients, such as the constant flow of detritus in lotic systems, used by detritivores. We opted for modelling the linear resources as it allows one parameter less than logistic growth, and it is also realistic*.

#### sort-organisms

Each organism draws a number, being sorted in descending order of the outcome. This number is randomly sampled from a normal distribution, with mean <u>energy</u>, and standard deviation <u>energy</u> * <u>stand-dev-to-body-size</u>.

*Rationale: This implementation gives advantage to organisms that have more <u>energy</u>, which have the opportunity to feed and reproduce first. Indirectly, this is advantageous to larger organisms, because they have more <u>energy</u> than smaller organisms at the same level of starvation. This implementation also allows smaller organisms with high <u>energy</u> (strong small organisms) to outcompete big organisms with low <u>energy</u> (weak large organisms). If <u>stand-dev-to-body-size</u> is very small (e.g*., *0.0001), the standard deviation parameter of the normal distribution becomes so small that essentially the sorting of organisms becomes deterministic. This is problematic under these settings as organisms evolve very quickly towards very large sizes, up to a point in which all die because their energy requirements are too high. On the contrary, if <u>stand-dev-to-body-size</u> is very large, the sorting is virtually random. With these settings, there is no advantage in having large <u>body-size</u>, and the simulation never stabilizes since organisms evolve towards becoming infinitely smaller*.

#### feeding

The energy intake of an organism (<u>energy-intake</u>) takes the minimum value of three possible quantities: <u>max-energy-intake</u>, their storage capacity (<u>max-energy</u> - <u>energy</u>) or the amount of resources in the patch they are on (<u>resources</u>). The <u>energy-intake</u> is then subtracted from the patch’s <u>resource</u>s.

*Rationale: Organisms will always try to feed as much as they can, limited by <u>max-resource-intake</u>. However, they might not be able to store all the incoming energy if their energetic income surpasses their <u>max-energy</u>. Additionally, they may only acquire as many <u>resources</u> as the environment allows (<u>resources</u>)*.

If the organism is not an alien, <u>energy-intake</u> is added to its <u>energy</u>.

*Rationale: This step distinguishes normal organisms from aliens. Alien organisms do not reproduce, nor die, but, like other organisms, they do take <u>resources</u> from patches. With aliens we want to create a lasting competition for resources. Aliens being immortal but without being able to reproduce ensures that both the native and alien populations are able to compete for the same <u>resources</u>. If both reproduce and die, then eventually one would exclude the other due to the competitive exclusion principle* (Hardin 1960).

#### movement

After **<u>feeding</u>**, the organism moves to another patch if the <u>energy-intake</u> is lower than <u>maintenance-cost</u> plus <u>maturation-cost</u>.

*Rationale: If an organism’s <u>energy-intake</u> is lower than the requirements for maintenance and maturation it might not be incorporating enough energy for its costs. In the case of adults, <u>maturation-cost</u> accounts for the accumulation of energy for reproduction*.

*If their <u>energy-intake</u> is not enough to cover those costs, organisms should be compelled to search for resources elsewhere. Otherwise there is no reason for the organism to move*.

The organism moves by selecting a random direction (<u>heading</u> is randomly assigned to an integer between 0 and 359, covering the entire spectrum of directions in a Netlogo model), and a random distance between 0 and <u>disp-ability</u>. If the organism is not an **alien**, they pay for the cost of movement, which is proportional to the distance dispersed (<u>energy</u> = <u>energy</u> - <u>dispersal-cost</u> * dispersal distance).

*Rationale: The dispersal algorithm models a simple form of dispersal, which is essentially a Brownian motion (Brown 1828), or isotropic random walk (Codling Edward A et al. 2008)*.

*Nonetheless, it is the form of choice of many organisms, like ballooning spiders and wind-dispersed plants*.

#### reproduction

If <u>age</u> is larger than <u>maturity-age</u> the organism is considered to be mature, and may reproduce (asexually). A mature organism may only reproduce if its <u>energy</u> exceeds <u>energy-to-reproduce</u>. The energy that is allocated to offspring is the difference between the <u>energy</u> level and <u>energy-after-reprod</u>. Then, each offspring receives an equal fraction of the allocated energy (equal to the allocated-energy divided by <u>fecundity</u>). Each offspring inherits the traits of the parent with a slight **<u>mutation</u>**, and its <u>age</u> set to 0.

*Rationale: We chose 0.3 and 0.7 for <u>energy-to-reproduce</u> and <u>energy-after-reprod</u> arbitrarily. These values allow an adult organism to be able to survive after reproduction, as their energy levels will be almost one third of the maximum. To reflect different life-forms, we ran robustness tests in which we increased the <u>energy-to-reproduce</u> (to reflect life-histories that only reproduce when their energy values are close to their maximum energy) and decreased the <u>energy-after-reprod</u> (organisms that barely survive after reproduction)*.

#### mutation

**<u>Mutation</u>** is called within the **<u>reproduction</u>** module. The offspring trait values (<u>body-size</u>, <u>maturity-age</u>, <u>fecundity</u>, and <u>disp-ability</u>) are sampled from a normal distribution with the mean parameter being the trait value of the parent, and spread parameter being the trait value of the parent multiplied by <u>mutation-amplitude</u>:

offspring-trait-value = N(parent-trait-value, parent-trait-value * mutation-amplitude)

*Rationale: <u>mutation-amplitude</u> controls the speed of mutation. We studied the impact of varying the mutation amplitude on robustness tests (Supplementary Materials S2)*.

For <u>fecundity</u> and <u>maturity-age</u>, an additional random variable is drawn from an uniform distribution between 0 and 1. If the value of this random value is smaller than the <u>offspring-trait-value</u> minus its floor integer the trait value is rounded to its ceiling value, and rounded to its floor value if otherwise.

*Rationale: <u>fecundity</u> and <u>maturity-age</u> are discrete traits, and therefore their values need to be rounded up or down, to either the next larger (ceiling) or smaller (floor) integer. Rounding to the nearest integer is not a good solution, because for small values of traits (e.g. 2) and typical values of mutation size (0.05) the probability of drawing a number from a normal distribution with mean 2 and standard deviation 0.05 * 2 that is below 1.5 or above 2.5 is very small*.

If <u>fecundity</u> or <u>maturity-age</u> are 0, they become 1.

*Rationale: Forcing <u>fecundity</u> and <u>maturity-age</u> to be at least 1 makes it so there are no organisms that reproduce zero offspring, or that are born mature already*.

#### maturation

If the **organism** is a juvenile, and not an **alien**, its <u>energy</u> is subtracted by <u>maturation-cost</u>.

*Rationale:* ***<u>maturation</u>*** *simulates costs of developing and assembling the body structures for organisms to be able to reproduce. In many agent-based models, the attaining of maturation comes with attaining a given size at maturation (Sibly et al. 2013). We did not implement actual growth in the model for the sake of simplicity, but instead organisms dissipate energy to mature. This is in some way similar to the simplest dynamic-energy budget organismal model, in which organisms attain reproductive maturation after dissipating enough energy into an abstract state variable called maturation (Kooijman 2000; Martin et al. 2012)*.

#### maintenance

The organism subtracts <u>maintenance-cost</u> from <u>energy</u>.

*Rationale: The maintenance costs correspond to costs of keeping the body functional, such as vital organ function and homeostasis*.

If the <u>energy</u> level is less than zero the organism dies of starvation.

*Rationale: The decision to let organisms pay their <u>maintenance-costs</u> before calling any other procedure was made because otherwise organisms evolve towards infinite fecundity, which was happening under high intensity of direct killing. If* ***<u>maintenance</u>*** *comes before other procedures, then <u>fecundity</u> levels above a certain threshold are not possible, otherwise newborn organisms die immediately*.

#### mortality

The organism draws a random number between 0 and its <u>age</u>. If the random number is higher than his <u>longevity</u>, it dies.

*Rationale: If the organism’s age is lower than <u>longevity</u>, it will never die before attaining an <u>age</u> at least equal to <u>longevity</u>. As <u>age</u> increases, the probability of sampling a number that is higher than <u>longevity</u> increases. Thus, the probability of dying increases as the organism ages*.

If <u>threat-phase?</u> is true, the organism draws a random number between 0 and 100. If the number is lower than <u>direct_killing</u>, it dies.

#### ageing

<u>age</u> is incremented by 1.

#### check stopping conditions

*Rationale: The algorithm reports true or false depending on whether the stopping conditions have been attained. The algorithm reports true if there are no more organisms alive, or if the number of organisms, as well as mean trait values, have not changed considerably for a considerable amount of timesteps*.

If the number of iterations until the next checkpoint (<u>next-checkpoint</u>) is larger than 0, **<u>check-stopping-conditions</u>** reports false. If <u>next-checkpoint</u> is 0, its value is updated under **<u>set-next-checkpoint</u>**, and then a snapshot of five values is taken: the number of (native) organisms, mean <u>body-size, maturity-age, fecundity</u>, and <u>disp-ability</u> of the current timestep. The snapshots for each value are stored in a stack (<u>nr-organisms, means-body-size, means-maturity-age, means-fecundity, means-disp-ability</u>). Each stack contains successive snapshots of each metric taken whenever <u>next-checkpoint</u> is 0. If the size of a stack is less or equal to nr-of-means, **<u>check-stopping-conditions</u>** reports false. If otherwise, the most recent value is compared with the mean of the previous (<u>nr-of-means</u>) values. If, for every metric, the most recent value is within the mean of the previous values +-<u>stopping-conditions-threshold</u> * mean of the previous values, then **<u>check-stopping-conditions</u>** reports true.

*Rationale: The core part of* ***<u>check-stopping-conditions</u>*** *only runs when a sufficient amount of timesteps have passed since the beginning of the simulation. Upon initialization of the model, <u>next-check</u> is initialized as 100, to ensure that the model does not advance to the next stage too early. The simulation is considered to have stabilized if the number of organisms and the mean trait values have not changed considerably in relation to previous timesteps. The algorithm reports false if there are still too few checkpoints to compare with, to avoid situations in which the simulations would stop too soon. To calibrate the algorithm’s parameter values (<u>nr-of-means, stopping-conditions-threshold</u>, and <u>stopping-conditions-interval</u>), we visually inspected different combinations of these parameters against the other model parameters with graphs within Netlogo (Fig. S1.2). We picked the values of <u>nr-of-means</u> = 5, <u>stopping-conditions-threshold</u> = 0.05, and <u>stopping-conditions-interval</u> = 50 because they ensured that the model would not trigger stopping conditions too soon or too late after stable conditions were attained*.

If **<u>check-stopping-conditions</u>** returns true, and <u>threat-phase?</u> is false, <u>threat-phase?</u> becomes true, and the simulation transitions to the perturbation phase. If **<u>check-stopping-conditions</u>** and <u>threat-phase?</u> is true, the simulation ends.

**Fig S1.2.**
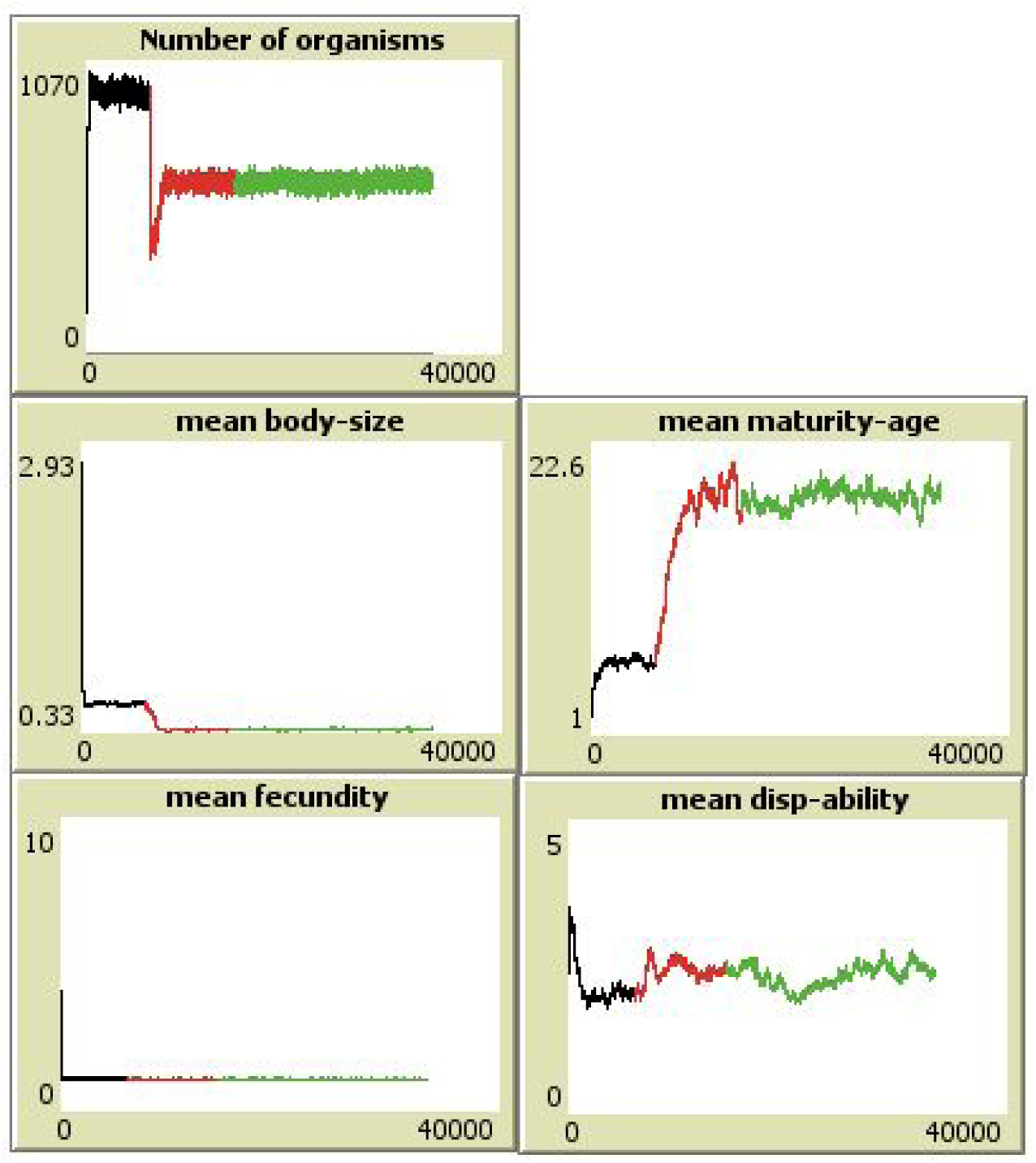
Plots showing the evolution of the number of organisms and mean trait values over time. Black line: before adding perturbation. Red line: stopping conditions were triggered and the perturbation (40% habitat degradation) is added. Green: Stopping conditions were triggered again, signaling the end of the perturbation phase. The graphs show that the triggering of stopping conditions happened at the right time, when the response variables have already attained stable values.

#### set-next-checkpoint

<u>next-checkpoint</u> is set to <u>stopping-conditions-interval</u> * mean of <u>age-at-reprod</u> of all organisms with age-at-reprod greater than 0.

*Rationale: The speed at which organisms attain maturity determines the speed at which organisms evolve. If organisms reproduce very quickly, the trait values quickly reach a new equilibrium. On the other hand, if organisms take longer to reach maturity then the convergence of trait values takes longer. In that case, the speed of change may be too slow to be detected between consecutive checkpoints of stopping conditions. Therefore, the interval between stopping conditions takes into consideration the mean time at reproduction of organisms*.

#### add-perturbations

##### Direct killing

Check **<u>mortality</u>** above.

*Rationale: Direct killing simulates background mortality rates*.

##### Habitat loss process

A random patch is selected and its <u>resources, resource-regen</u>, and <u>max-resources</u> set to 0. Then, neighbouring patches also have their state variables set to 0, until the maximum number of patches affected reaches the maximum, which is the total number of patches * <u>habitat_loss</u> / 100.

*Rationale: The growth of affected patches follows the same algorithm as the growth of resource patches around initially marked resource patches. Habitat loss simulates the loss of suitable habitat aggregated in space*.

##### Habitat fragmentation process

The <u>resources, resource-regen</u>, and <u>max-resources</u> of a random number of patches is set to 0. The number of random patches selected is equal to total number of patches * (<u>habitat_fragmentation</u> / 100).

*Rationale: Habitat fragmentation simulates the reduction in suitable habitat, when the reduction of habitat happens randomly in the landscape*.

##### Habitat degradation process

The <u>resources, resource-regen</u>, and <u>max-resources</u> of all patches is multiplied by (1 - <u>habitat_degradation</u>).

*Rationale: Habitat degradation simulates the reduction of resource availability*.

##### Invasion process

A random number of organisms creates a copy of themselves (**aliens**), inheriting all their state variables. Aliens do not reproduce (check **<u>reproduction</u>**), do not gain or lose <u>energy</u> (check **<u>feeding, movement, maturation</u>**, and **<u>maintenance</u>**), and do not die. The number of aliens added is equal to number of organisms * (<u>invasives</u> / 100).

*Rationale: In the real world, alien invasive species often are successful in relation to the natives because they have no antagonists (predators, diseases, parasites, etc.). Our invasive organisms simulate an invader competing for the same resources as the native organisms. We simulate invasive pressure by adding new organisms to the model that are functionally similar to the natives. To ensure co-existence of aliens and natives, we implemented aliens as immortal organisms, without the capacity to reproduce. This guarantees competitive pressure for resources*.

**Table S1.3:** Model parameters, their description, their permitted range and values used for the standard simulations and robustness tests.

| Parameter name | Description | Range | **Standard**, and tested values |
| --- | --- | --- | --- |
| Landscape and initialization |  |  |  |
| <u>w-width</u> | Width and length of landscape | 1-∞ [int.] | **33**, 11, 99 |
| <u>w-length</u> |  |  |  |
| <u>nr-resource-clusters</u> | Randomness of resource distribution | 1-544 [int.] | **33**, 11, 99 |
| <u>resource-patch-fract</u><br><u>ion</u> | Proportion of resource patches | 0-1 | **0.5**, 0.16, 1 |
| <u>map-seed</u> | Random seed to replicate maps | 0-100 | **0**, - |
| <u>starting-nr</u> | starting number of organisms | 1-∞ [int] | **500** |
| Patches |  |  |  |
| <u>max-resources</u> | Maximum resources | 1 [const.] | **1** |
| <u>regen-rate</u> | Regeneration rate of resources | 0-1 | **0.2**, 0.06, 1 |
| Energy budgets |  |  |  |
| <u>max-energy-intake-coef</u> | intake and expenditure relative to the maximum energy of organisms | 0-1 | **0.3**, -,0.9 |
| <u>maintenance-cost-coef</u> |  | 0-1 | **0.1**, 0.03, 0.2 |
| <u>maturation-cost-coef</u> |  | 0-1 | **0.1**, 0.03, 0.2 |
| <u>dispersal-cost-coef</u> |  | 0-1 | **0.01**, 0.003, 0.03 |
| <u>energy-to-reproduce-coef</u> | Minimum amount of energy needed to reproduce and energy after reproduction, relative to maximum energy of organisms | 0-1 | **0.7**, -, 0.9 |
| <u>energy-after-reprod-coef</u> |  | 0-1 | **0.3**, 0.1, 0.9 |
| Other |  |  |  |
| <u>metabolic-allometric-exponent</u> | Influences energetic intakes and expenditure of energy in relation to body size | 0.66-1 (typical range across the tree of life) | **0.75**, 0.66, 1 |
| <u>maturity-longevity-coef</u> | Dependence of longevity on maturity | 0-∞ | 3, 1, 9 |
| <u>stand-dev-to-body-size</u> | Strength of energy sorting | 0-∞ | 1, -, - |
| Mutation |  |  |  |
| <u>mutation-amplitude</u> | Degree to which mutation occurs | 0-∞ | 0.05, 0.016, 0.15 |
| <u>body-size-mutation?</u> | Activates mutation for these traits | true, false<br>[boolean] | true |
| <u>maturity-age-mutation?</u> |  | true, false<br>[boolean] | true |
| <u>fecundity-mutation?</u> |  | true, false<br>[boolean] | true |
| <u>disp-ability-mutation?</u> |  | true, false<br>[boolean] | true |
| Stopping conditions |  |  |  |
| <u>stopping-conditions-threshold</u> | Allowed deviation from previous checkpoints | 0-∞ [ratio] | 0.05 |
| <u>stopping-conditions-interval</u> | Number of organism generations until next checkpoint | 1-∞ [int.] | 50 |
| <u>nr-of-means</u> | Number of previous checkpoints to compare with | 1-∞ [int] | **5** |
| Perturbations |  |  |  |
| <u>direct_killing</u> | random killing of organisms each timestep | 0-100 [%] | **0-10** |
| <u>habitat_loss</u> | aggregated destruction of suitable habitat patches | 0-100 [%] | **0-80** |
| <u>habitat_fragmentation</u> | random destruction of habitat patches | 0-100 [%] | **0-70** |
| <u>habitat_degradation</u> | decrease in habitat quality of all patches | 0-100 [%] | **0-50** |
| <u>invasives</u> | introduction of immortal organisms | 0-100 [%] | **0-70** |
| Burn-in recycling |  |  |  |
| <u>filename</u> | file in which to save finished burned-in simulation | [string] | - |
| <u>only-run-burn-in?</u> | whether to run only<br>burn-in (and save it to<br>disk) | true/false<br>[boolean] | **true/false** |
| <u>load-world?</u> | loads an already<br>“burn-ed in” simulation | true/false<br>[boolean] | **true/false** |
| Aesthetics |  |  |  |
| <u>turtle-aesthetics?</u> | size of organisms as<br>appears in the Netlogo<br>interface is proportional<br>to body size | true/false<br>[boolean] | **false** |

## Supplementary Materials S2: Robustness analyses

The Robustness analyses test whether varying the parameter values impacts the results of the experiment. Parameter values were selected so that they would reflect extreme conditions (Table S1.3). In most cases, extreme parameter values were selected as being three times lower and higher than standard trait values (thus capturing about a magnitude change in trait values). For parameters that are bounded on one side (e.g. energetic costs, resource clusters), we either selected the extreme low and high values for that parameter (maximum resource patch fraction is 1), or we selected a value close to the extreme (maintenance costs cannot be higher than 0.2, otherwise the energy intake of organisms is lower than the energy expenses). In some cases, selecting either an extreme low or extreme high did not make sense as the standard value was already in such extreme low or high value (relative energy intake, energy to reproduce, energy-after-reprod). We varied the parameters locally, i.e., varying only one parameter at a time. In the case of the metabolic allometric exponent, we only tested the typical values found in the literature: 0.66 to 1.

**Table S2.1:**
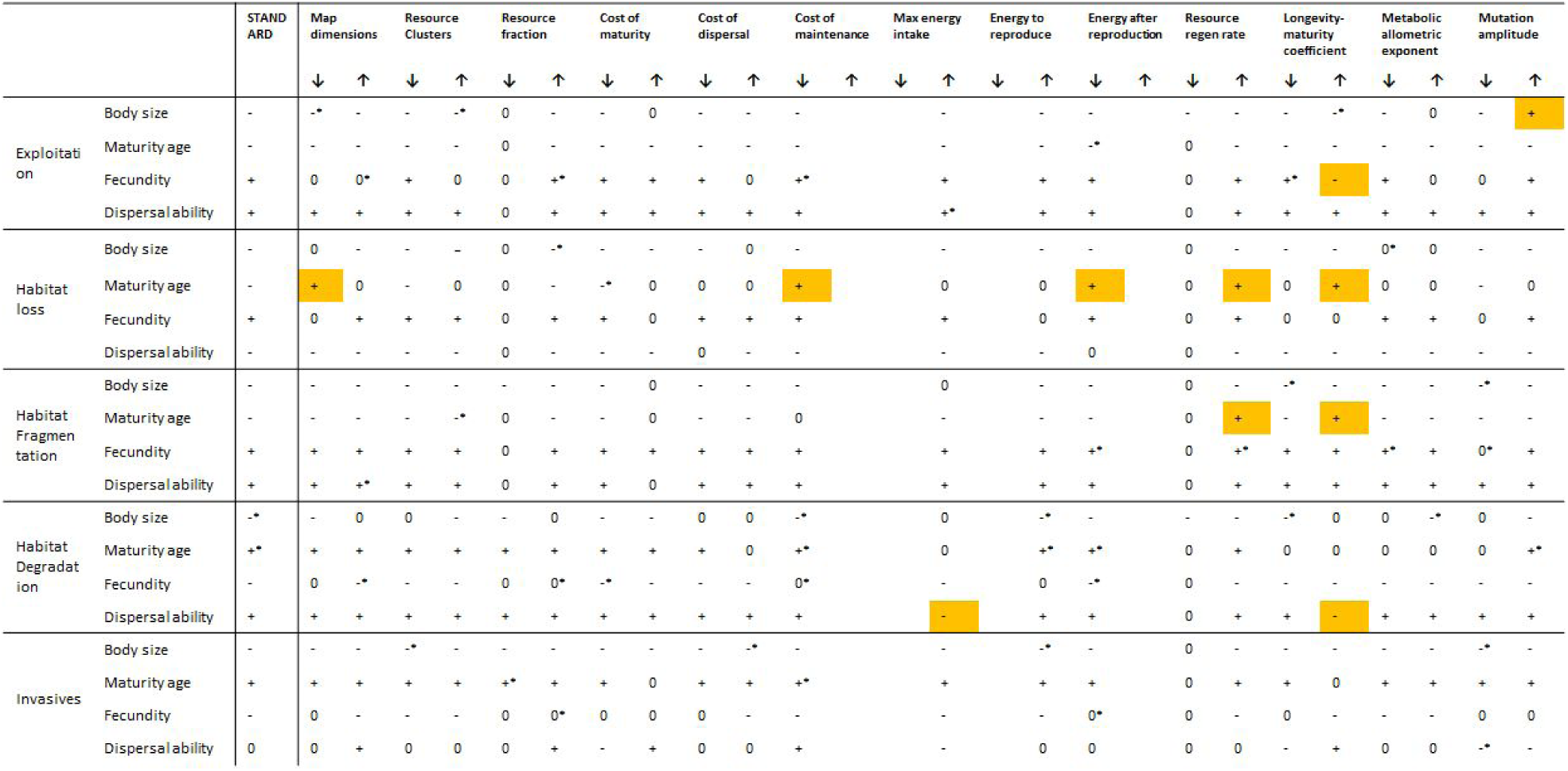
Direction of the relative change in trait values for each robustness analysis. “ +” indicates positive change in trait value. “ -” Indicates negative change. “ *” indicates cases in which the signal was inspected visually, as the GLMM did not converge. In orange, cases where the signal is opposite to that of the standard runs.

## Supplementary Materials S3: Data and scripts

Supplied as separate files:

- Netlogo Model: **Logotraits19.nlogo**
- R scripts to run Netlogo through R: **Logotraits_main.R** and **Logotraits_interface.R**
- R scripts to run Statistical analyses: **Data_analysis.R**
- Datasets of the standard and robustness analyses, respectively: **SA_dataset.csv** and **RA_dataset.csv**

